# DEAR-OWL: a fully browser-based hybrid resource for instant or precise differential gene expression analysis

**DOI:** 10.64898/2026.07.28.741369

**Authors:** Kota Kambara, Sintho Wahyuning Ardie, Daisuke Tsugama

## Abstract

**Motivation:** Differential gene expression analysis (DEA) via RNA sequencing (RNA-seq) is essential but remains challenging for wet-lab biologists due to command-line complexities. Centralized web platforms democratize this process but suffer from server congestion, long queuing delays, data privacy risks with proprietary datasets, and limited long-term sustainability due to hosting fees.

**Results:** We present DEAR-OWL (Differential Expression Analysis Resource on the Web (Lite)), a fully serverless, privacy-preserving web application that performs the DEA locally inside the user’s web browser. To combine instant exploratory speed with rigorous verification, the application runs two distinct analysis options. The first option is a fast screening tool written in native browser language (JavaScript) that delivers immediate, genome-wide fold-change calculations and statistical screening based on an edgeR-equivalent logic. The second option is a heavy-duty statistical tool that brings the standard R package (DESeq2) directly into the browser using WebR and WebAssembly technology, ensuring publication-grade validation without needing server power. Interactive visual plots (volcano plots, minus-average plots, and heatmaps) are seamlessly generated from the results of either analysis choice. Benchmarking proved its hardware compatibility: the browser-based DESeq2 engine completed the analysis in ∼30 seconds on a 64 GB RAM workstation and in ∼3 minutes on an 8 GB RAM laptop without crashing. DEAR-OWL can utilize the Grass Expression Atlas (GExA) data as built-in and supports secure local file uploads, ensuring total data privacy with neither queuing delays nor cloud infrastructure costs.

**Availability and implementation:** DEAR-OWL is freely accessible at https://webpark2116.sakura.ne.jp/deseq2/. The source code is available at https://github.com/kota200/DEAR-OWL.

## Introduction

Differential gene expression analysis (DEA) using RNA-sequencing (RNA-seq) data has become an indispensable workflow in modern biomedical and agricultural research. Standard bioconductor packages in R, such as DESeq2 (Love *et al*. 2014) and edgeR (Robinson *et al*. 2010), are widely recognized as the gold standards for statistical rigor in transcriptomic quantification. Despite their ubiquity, executing these CLI (Command Line Interface)-based tools remains a formidable bottleneck for wet-lab biologists who lack formal training in programming, package dependency management, and/or computing environments.

To bridge this gap, web-based graphical user interfaces (GUIs), including Galaxy (Galaxy Community 2022) and various R-Shiny (Chang *et al*. 2026) applications, have been developed to democratize transcriptomic analysis. However, these conventional web platforms inherently rely on a centralized, server-side paradigm, which introduces three limitations. First, centralized servers can suffer from infrastructure congestion; high concurrent user volumes lead to extensive queuing delays, stretching analysis turnaround times from minutes to hours or even days. Second, uploading large genomic count matrices or raw clinical datasets to third-party external servers raises data privacy and security concerns, particularly in medical and proprietary industrial research. Third, the long-term sustainability of these web servers is perpetually constrained by hosting fees and maintenance overhead, frequently resulting in abandoned or broken web links over time.

Recent advancements in WebAssembly (Rossberg 2019) and the emergence of webR (Stagg and Lionel 2023), which can compile the R environment directly into the web browser, offer a paradigm shift toward decentralized, client-side bioinformatics. By leveraging the computational resources of the user’s local machine, client-side execution can theoretically eliminate server dependency entirely while preserving maximum data privacy.

Here, we present a novel web application, DEAR-OWL (DEA Resource On the Web (Lite)) designed for instant DEA. To combine instant exploratory speed with rigorous verification, the application runs two distinct analysis options. The first option is a fast screening tool written in native browser language (JavaScript) that delivers immediate, genome-wide fold-change calculations and statistical screening based on an edgeR-equivalent logic. The second option is a heavy-duty statistical tool that brings the standard R package (DESeq2) directly into the browser using WebR technology, ensuring publication-grade validation without needing server power. Because all computations are executed within the local sandbox environment, data never leaves the user’s computer, ensuring privacy with neither queuing delays nor cloud infrastructure costs. This application redefines the user experience of DEA, providing a seamless, secure, and instantaneous pipeline from raw count matrices to publication-ready data visualization.

## Materials and Methods

### System architecture and implementation

DEAR-OWL was constructed using standard frontend web technologies, primarily HTML5 and native JavaScript (ES6+), ensuring high portability and cross-platform compatibility without requiring any server-side installations. The core statistical workflow relies on webR (v0.2.2 or later), which compiles the R statistical environment into WebAssembly. This allows the application to execute standard Bioconductor packages, specifically DESeq2, directly within the browser’s local sandbox environment. To prevent UI freezing during heavy statistical computations, the webR runtime and data processing pipelines are offloaded to an isolated Web Worker thread.

For the fast screening pipeline, we implemented a custom, lightweight differential expression engine written entirely in native JavaScript. To replicate the statistical properties of traditional count-based frameworks, this engine calculates log_2_ Counts Per Million (log_2_CPM) for each gene using an edgeR-equivalent dynamic prior-count scaling method. The mathematical pipeline is executed as follows:

1. **Dynamic Prior-Count Scaling**: For each sample *s*, a dynamic prior count (adjPrior_*s*_) is derived from a baseline prior count (prior.count = 2.0) scaled by the ratio of the sample’s library size (*L*_*s*_) to the average library size across all samples (avgLibSize), such that adjPrior_*s*_ = 2.0 × (*L*_*s*_/avgLibSize).
2. **log2CPM Transformation**: The raw count *C*_*g,s*_ for gene *g* in sample *s* is transformed into a stabilized log_2_CPM value via the formula:

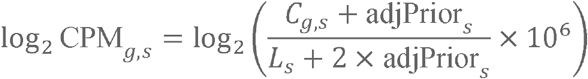
3. **Statistical Testing via Z-score**: The log_2_ fold change (LFC) is computed as the difference between the mean log_2_CPM values of the treatment and control groups. To assess statistical significance across replicates, a two-tailed Z-test is performed. The standard error is calculated with a variance-stabilizing lower-bound threshold of 0.01 to prevent inflation from zero-variance artifacts. The definitive *p*-value is then derived using a high-precision Gaussian error function (erf).
4. **Multiple Testing Correction**: The resulting raw *p*-values are dynamically adjusted for multiple comparisons using either the Benjamini-Hochberg false discovery rate (FDR) (Benjamini and Hochberg 1995) or the Bonferroni (1936) correction method, according to the user’s configuration.

### Data integration and input modalities

The platform provides two distinct modalities for data input. First, the platform features integration with the Grass Expression Atlas (GExA) (Kambara *et al*. 2026), allowing users to import pre-processed public RNA-seq datasets (covering thousands of samples from *Pennisetum glaucum, Setaria italica*, etc.) for immediate analysis. Second, the interface supports custom user uploads of raw gene count matrices in a CSV or TSV format. All uploaded data are processed client-side, and no datasets are transmitted to external servers.

## Results and Discussion

### Hybrid dual-engine architecture and interface

Our platform features a single-page graphical user interface (GUI) for DEA. Users can either load the built-in GExA data or upload a custom raw gene count matrix directly via the browser interface. To balance analytical speed with statistical rigor, the platform offers a dual-engine workflow. For rapid screening, the lightweight native JavaScript engine executes an edgeR-like test. Alternatively, for comprehensive statistical validation, the client-side DESeq2 engine initializes a webR instance inside an isolated Web Worker thread, downloads the required Bioconductor packages dynamically, and executes the complete, unaltered DESeq2 pipeline—including median-of-ratios normalization, dispersion estimation, and statistical testing with various configurable options—entirely within the isolated local environment. Following the execution of either engine, interactive volcano plots, minus-average (MA) plots, and sample correlation heatmaps are instantly updated in real time (Figure 1).

**Figure 1.**
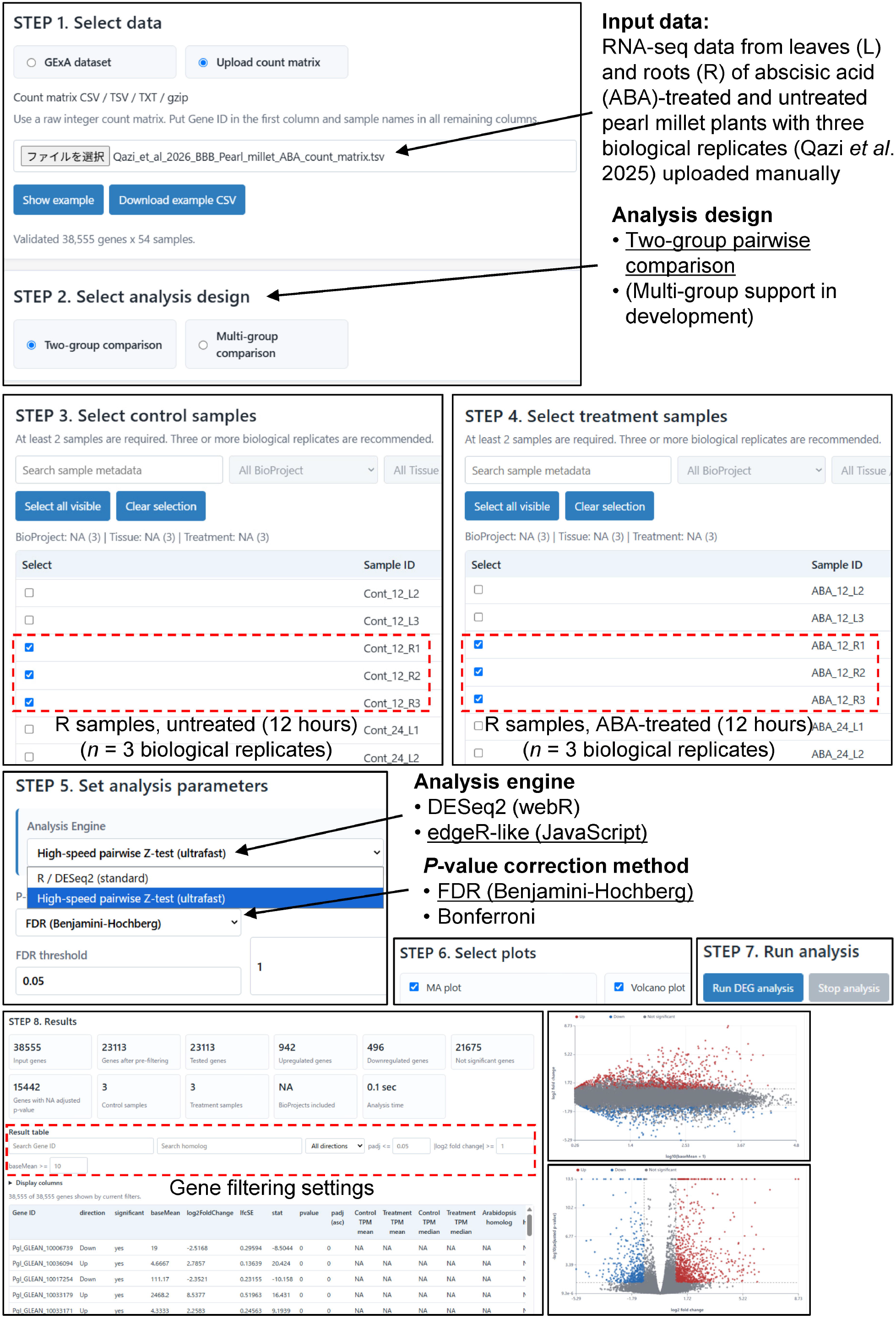
Example of DEA using DEAR-OWL. The figure exemplifies a two-group pairwise comparison using a manually uploaded RNA-seq dataset from leaves (L) and roots (R) of pearl millet plants (*Cenchrus americanus*) treated with or without abscisic acid (ABA), utilizing three biological replicates (Qazi *et al*. 2025). Available options for the analysis design (for Step 2), execution engine (Step 5), and *P*-value adjustment (Step 5) are explicitly labeled, with the currently selected options (Two-group pairwise comparison, the custom edgeR-like (JavaScript) engine, and the FDR (Benjamini– Hochberg) correction method, respectively) indicated by underlines.

### Computational performance and benchmarking

To evaluate the practical feasibility of our web application, we benchmarked the execution times using a real-world *Pennisetum glaucum* (pearl millet) RNA-seq dataset (Qazi *et al*. 2025), two computational environments, and Google Chrome. On a workstation equipped with 64 GB of RAM, the client-side DESeq2 pipeline via webR completed the entire analysis in approximately 30 seconds. On a laptop with 8 GB of RAM, the same pipeline completed the analysis in approximately 3 minutes without triggering any browser out-of-memory (OOM) crashes. While the performance variance likely stems from differences in CPU processing power and browser-level memory allocation, these results demonstrate that our platform is fully operational on standard hardware.

This decentralized, client-side paradigm directly addresses the long-standing infrastructure challenges faced by conventional centralized web servers, such as server queuing delays and data privacy risks. By eliminating server-side computation entirely, our platform ensures permanent availability with zero long-term maintenance overhead. Although highly complex datasets with hundreds of samples may still benefit from high-performance computing clusters, our findings confirm that for standard experimental designs, browser-based execution provides a practical, secure, and instantaneous alternative for wet-lab biologists.

## Acknowledgements

The authors appreciate the computational resources provided by The University of Tokyo.

## Funding

This work was supported by Japan Society for the Promotion of Science (JSPS) KAKENHI grants 25K09059 (to D.T.) and 24KJ0912 (to K.K.).

## Declaration on the use of generative AI

During the preparation of this manuscript and the development of the tool, the authors utilized a large language model (Gemini 1.5 Flash, Google) to generate the initial structural outlines, technical drafts of the source code, and documentation, with extensive assistance particularly for the frontend user interface implementation and writing adjustments to improve text clarity. Following this AI-assisted phase, the authors rigorously reviewed, edited, and verified all code components and text to ensure scientific accuracy, technical validity, and compliance with academic standards, taking full responsibility for the final content and the developed tool.

## Code availability

The source code for DEAR-OWL is open-source and publicly available on GitHub at https://github.com/kota200/DEAR-OWL.

## Data availability

The built-in public RNA-seq datasets featured in DEAR-OWL are integrated from the Grass Expression Atlas (GExA). These datasets can be reproduced and verified using the biological accession numbers and metadata structures described in the original GExA publication (Kambara *et al*. 2026).

## Author contributions

D.T. and K.K. conceived the study. K.K., S.W.A. and D.T. implemented the platform. D.T. wrote the original draft, and K.K., S.W.A. and D.T. reviewed and edited the manuscript. D.T. and K.K. acquired the funding for this study.

## Competing interests

The authors declare no competing interests.

## Notes

https://github.com/kota200/DEAR-OWL

https://webpark2116.sakura.ne.jp/deseq2/

### Competing Interest Statement

The authors have declared no competing interest.

